# An Efficient Computational Method to Create Positive NIPT Samples with Autosomal Trisomy

**DOI:** 10.1101/2023.11.24.568620

**Authors:** Tinh H. Nguyen, Huynh V. Nguyen, Minh D. Pham, Vinh S. Le

**Affiliations:** University of Engineering and Technology, Vietnam National University, Ha Noi, Viet Nam; GENTIS, Hanoi, Vietnam

**Keywords:** NIPT, data simulation, mosaicism, low coverage data, autosomal chromosome aneuploidy

## Abstract

Noninvasive prenatal test (NIPT) has been widely used for screening trisomy on chromosomes 13 (T13), 18 (T18), and 21 (T21). However, the false negative rate of NIPT algorithms has not been thoroughly evaluated due to the lack of positive samples. In this study, we present an efficient computational approach to create positive samples with autosomal trisomy from negative samples. We applied the approach to establish a low coverage dataset of 1440 positive samples with T13, T18, and T21 aberrations for both mosaic and non-mosaic conditions. We examined the performance of WisecondorX and its improvement, called VINIPT, on both negative and positive datasets. Experiments showed that WisecondorX and VINIPT with a z-score threshold of 3.3 were able to detect all non-mosaic samples with T13, T18, and T21 aberrations (i.e., the sensitivity of 100%). Using a lower z-score threshold of 2.58 when analyzing mosaic samples, both WisecondorX and VINIPT have the overall sensitivity of 99.7% on detecting T13, T18, and T21 aberrations from mosaic samples. WisecondorX has the specificity of 98.5% for the non-mosaic analysis, but a considerably lower specificity of 95% for the mosaic analysis. VINIPT has a much better specificity than WisecondorX, i.e., 99.9% for the non-mosaic analysis, and 98.2% for the mosaic analysis. The results suggest that VINIPT can play as a powerful NIPT tool for the low coverage data.

## I. Introduction

The abnormality in the number of chromosomes results in different disorders in both males and females. The well-known chromosome abnormalities are the trisomy 13 (T13 or Patau syndrome), trisomy 18 (T18 or Edwards syndrome), and trisomy 21 (T21 or Down syndrome). The NIPT using the cell-free DNA (cfDNA) which is a mixture of maternal DNA and a small percentage of fetal DNA in the blood of mother has been widely applied to screen the autosomal disorders [1]–[6]. We note that the cfDNA from mother and fetus are mixed and not feasibly separated.

A number of computational algorithms have been developed to predict chromosome aneuploid based on whole genome data sequenced from cfDNA such as NIFTY [7], NIPTeR [8], Wisecondor [9], WisecondorX [10], and triSure [11]. Experiments on a limited number of clinically validated positive samples showed that the NIPT algorithms have high sensitivity and specificity on screening T13, T18, and T21 [6], [12], [13], e.g., the sensitivity of 100% for T13, 98.2% for T18, and 99.1% for T21 [6]; and the specificity of around 99.9% for determining negative samples.

The specificity of the NIPT algorithms can be measured from large-scale and timely retrospective studies. However, the sensitivity (or the false negative rate) of the NIPT algorithms, one of the most important indicators, is not easily measured due to the lack of positive samples. Up to date, the positive datasets collected from hospitals or genetic testing centers are normally small and not available for scientific community. A large number of negative (normal) NIPT samples can be gathered from the hospitals or genetic testing centers. We note that the performance of NIPT with mosaic samples has not been systematically evaluated.

Simulation can play as a reasonable alternative way to examine the sensitivity of NIPT algorithms. Although a number of simulation methods have been proposed to generate general genomic datasets [14], no simulation method has been specifically designed to simulate NIPT data. The genomic data obtained from NIPT come mostly from the genome of mother and only a small fraction (normally around 10%) from the fetal genome. Generating short-reads for NIPT data is very challenging due to the complicated genomic characteristics of the NIPT data such as DNA artifacts, fragment size distribution, short-read quality and distribution, sequencing errors, or GC-content variation.

In this paper, we present a computational method to create positive samples with autosomal trisomy from negative samples. Instead of generating and adding new short-reads into the chromosome of interest *h*, we will remove short-reads of the other chromosomes such that the read coverage of the chromosome *h i*n comparison with that of the other chromosomes indicates three copies of chromosome *h* in the fetal genome. We describe how to create positive samples from negative samples for both non-mosaic and mosaic conditions. The generated positive datasets were used to measure the sensitivity of the NIPT algorithms. We also collected a dataset of 1058 negative samples to measure the specificity of the NIPT algorithms.

## II. Methods

We collected a dataset of 1500 samples from singleton pregnancies with clear negative predictions from NIPT and no customer reports about false results. The NIPT samples were sequenced by the MGISEQ-200 platform with 50 bp single-end reads. The samples with the minimum of 5 million reads were selected for the study.

The short-reads were aligned with the reference genome GRCh37 by Bowtie 2 [15] to create alignments. We employed the reference GRCh37 instead of the reference GRCh38 because some NIPT programs use pretrained models that are not available for the reference GRCh38. The reads that mapped to repeat or centromeric regions were removed to avoid data noises.

Let *C*_*i*_ be the read percentage (read coverage) mapped to chromosome *i*, e.g., *C*_1_ = 0.08 means that the number of reads in chromosome 1 counts for 8% of all reads in the whole genome data. The reads mainly come from the DNA of mother and a small fraction from the DNA of fetus. Let *f* be the fetal DNA fraction of the sample, i.e., the percentage of reads from the fetus. For example, *f* = 9% indicates that 9% of reads come from the genome of fetus, and the remaining 91% of reads come from the mother’s genome. The fetal DNA fraction *f* can be estimated by some computational methods of which SeqFF [16] can reliably estimate the fetal DNA fraction for both male and female NIPT samples. The fetal DNA fraction *f* might be overestimated, especially for samples with a low fetal DNA fraction. To avoid creating easy positive samples that might make NIPT algorithms overconfident about their sensitivity, we decrease the value of *f* to its 90% (i.e., *f* = 0.9 × *f*) in the study. As a result, the generated positive samples might be slightly more difficult than the real positive datasets.

We divided the 1500 negative samples into two datasets: a dataset **P** of 1200 samples for evaluating the specificity of NIPT algorithms and a dataset **S** of 300 samples used to create positive samples. We removed all samples with small fetal DNA fractions (i.e., *f* smaller than 5%). As a result, the dataset **P** consists of 1058 samples and the dataset **S** contains 240 samples.

### A. Generating positive non-mosaic samples with autosomal trisomy

A normal sample *s* has two copies of every autosomal chromosome. The sample *s* has a trisomy at a chromosome *h* if the fetus has three copies of *h*. Given a negative sample, instead of adding new reads to create an additional copy of *h* for the fetus, we remove available reads in the other chromosomes such that the read percentage of *h* in comparison with that of the other chromosomes will be equivalent to the case that the fetus has three copies of *h*. Technically, for a singleton non-mosaic sample, introducing an additional copy of *h* for the fetus will increase the read percentage *C*_*h*_ to *C*′_*h*_ = *C*_*h*_(1 + 0.5 × *f*). Increasing the read percentage *C*_*h*_ to *C*′_*h*_ is equivalent to decreasing the read coverage *C*_*i*_ of the other chromosome 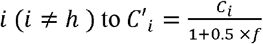.

### B. enerating positive mosaic samples with autosomal trisomy

We also generated positive mosaic samples. There are different types of mosaicism that strongly affect the accuracy of the NIPT methods [17]. In this study, we generated a common type of mosaic case (placenta mosaicism) that half of fetal cells are normal with 46 chromosomes and the other half are abnormal with the autosomal trisomy. The read percentage *C*_*h*_ of the mosaic sample will be increased to *C* ′′_*h*_ = *C*_*h*_(1 + 0.25 × *f*). In the mosaic case, the difference between *C*′′_*h*_ and *C*_*h*_ is only half of the difference between *C*′_*h*_ and *C*_*h*_. Thus, detecting aberrations in the mosaic samples is much more difficult than that in the non-mosaic samples. Given a negative sample *s* and chromosome *h*, for every chromosome *i* (*i* ≠ *h*), we remove reads such that the read percentage *C*_*i*_ will be reduced to 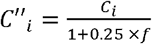.

We generated positive samples from 240 negative samples under both non-mosaic and mosaic conditions. We obtained a positive dataset of a total of 1440 positive samples with T13, T18, and T21 aberrations for testing the NIPT algorithms.

### C. NIPT algorithms

Up to date, WisecondorX [10], an improvement of the Wisecondor algorithm [9], is the one of the best open-source NIPT algorithms for the low coverage NIPT data [13]. The typical workflow of the read coverage-based NIPT algorithms include four steps:

1. Short reads are aligned with the reference genome. Artifact duplications, ambiguous mapped reads, or reads mapped into repeat regions should be eliminated.
2. The genome is divided into a serial of equally sized bins. The number of reads for every bin will be counted from the alignment. The GC-correction technique or any method to correct the read coverage bias might be applied to reduce data noises. For the low coverage data, we should use a large bin size (at least 10^5^ bps) such that the number of reads per bin is sufficient.
3. Given a testing sample *s*, its read coverages will be analyzed with that of normal samples through statistical testing. The statistical test will figure out bins on *s* whose read coverage significantly deviates from the expected read coverage of a normal sample. To avoid the read coverage variation across different samples, the within-sample analysis methods such as Wisecondor/WisecondorX compare a bin *b* on sample *s* to a set of reference bins on the other chromosomes within *s*. The bin *b* and its reference bins have similar behaviors and equivalent characteristics.
4. The bin aberrations of on chromosome *h* are combined to calculate the overall aberration score (i.e., z-score) of *h*. If the z-score of *h* is greater (or smaller) than a predefined z-score threshold, the chromosome *h* is considered to be abnormal. Otherwise, it is considered to be normal. The Wisecondor uses the Stouffer’s z-score sliding window approach to segment and determine aberrations. WisecondorX uses the circular binary segmentation algorithm [18] instead of Stouffer’s technique to detect segment aberrations that overcomes the running time burden of the Wisecondor algorithm.

We used WisecondorX to analyze our low coverage NIPT data and found a number of limitations. The first limitation of WisecondorX is a considerable number of fall positive samples when using the overall z-score to predict chromosome aberrations. The second limitation of WisecondorX is the sensitivity to reference samples used to create the reference panel. We developed a software of the so-called VINIPT, an improvement and adaption of Wisecondor and WisecondorX algorithms, optimized for the low coverage data. First, we determined proper reference samples to build a number of reference panels that help consistently predict the chromosome aberrations. Second, the VINIPT algorithm combines different algorithms to determine if a chromosome is abnormal. The improvements help reduce the false positive rate of the VINIPT algorithm.

We note that some NIPT algorithms such as NIFTY or triSure were implemented in commercial software and not publicly available for testing. Besides, WisecondorX and VINIPT algorithms, we also examined the open-source NIPTeR algorithm [8]. However, preliminary results showed that NIPTeR did not work well on our low coverage data. The performance of NIPTeR should be further investigated, therefore, we eliminated its results from the report.

## III. Results

In this paper, we reported the performance of WisecondorX and VINIPT software on both 1058 negative samples and 1440 generated positive samples. Fig. 1 shows the distribution of fetal DNA fractions and read coverages of the samples. The samples have the fetal DNA fractions ranging from 5% to 22% and the read coverages from 5M reads (i.e., about 0.08*x*) to 25M reads (i.e., about 0.4*x*). Most of the samples have fetal DNA fractions from 8 to 12 (i.e., the mean fetal DNA fraction of 9.45) and the read coverages from 9M reads to 15M reads (i.e., the mean read coverage of 11.8M).

**Fig 1.**
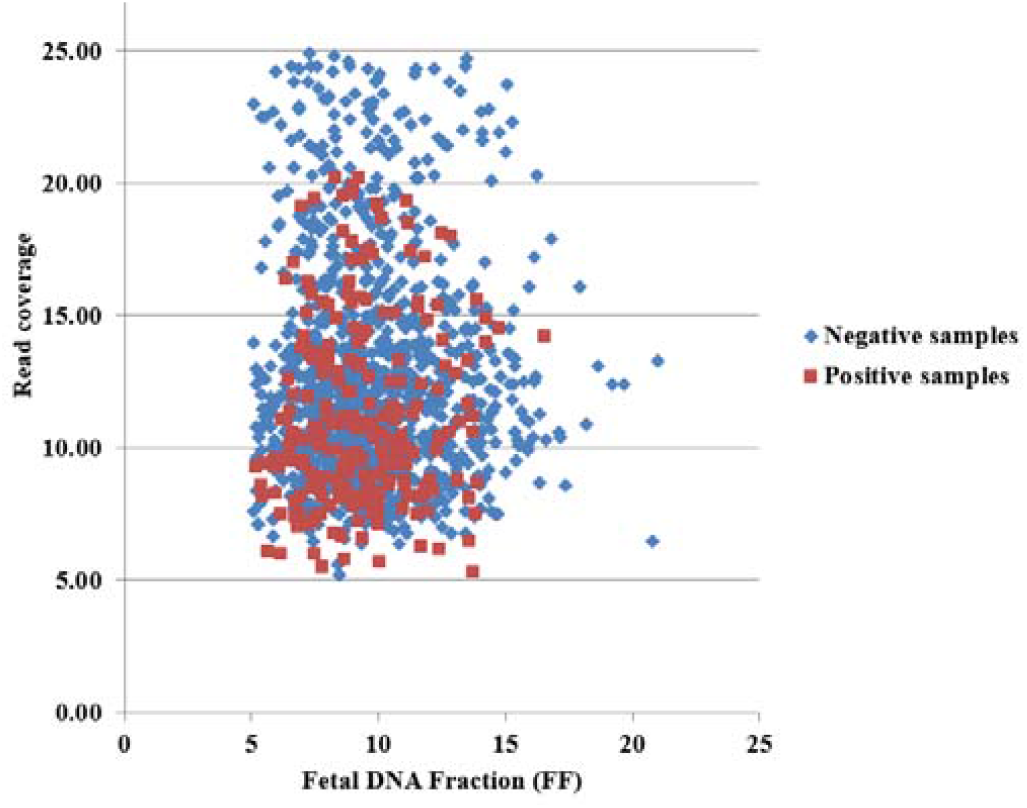
The distribution of fetal DNA fractions and read coverages of negative and generated positive samples

We analyzed the results from WisecondorX and VINIPT on negative samples and positive samples from non-mosaic samples (see Fig. 2). For the non-mosaic analysis, we set the z-score threshold of 3.3 for both WisecondorX and VINIPT. The z-scores of negative samples form one group and most of them smaller than 3.3; while the z-scores of positive samples form another separated group and most of them are greater than 3.3. Both WisecondorX and VINIPT could determine all T13, T18, and T21 aberrations on the positive samples, i.e., the sensitivity of 100%. The WisecondorX resulted in 16 false positive samples (i.e., 16 negatives samples were predicted as positive samples). The false positive rate of WisecondorX for non-mosaic samples is about 1.5% or the specificity of about 98.5%. The VINIPT falsely assigned only one negative sample as a positive sample with T13. The false positive rate of VINIPT is smaller than 0.1% (i.e., specificity greater than 99.9%) and much less than that of WisecondorX.

**Fig 2.**
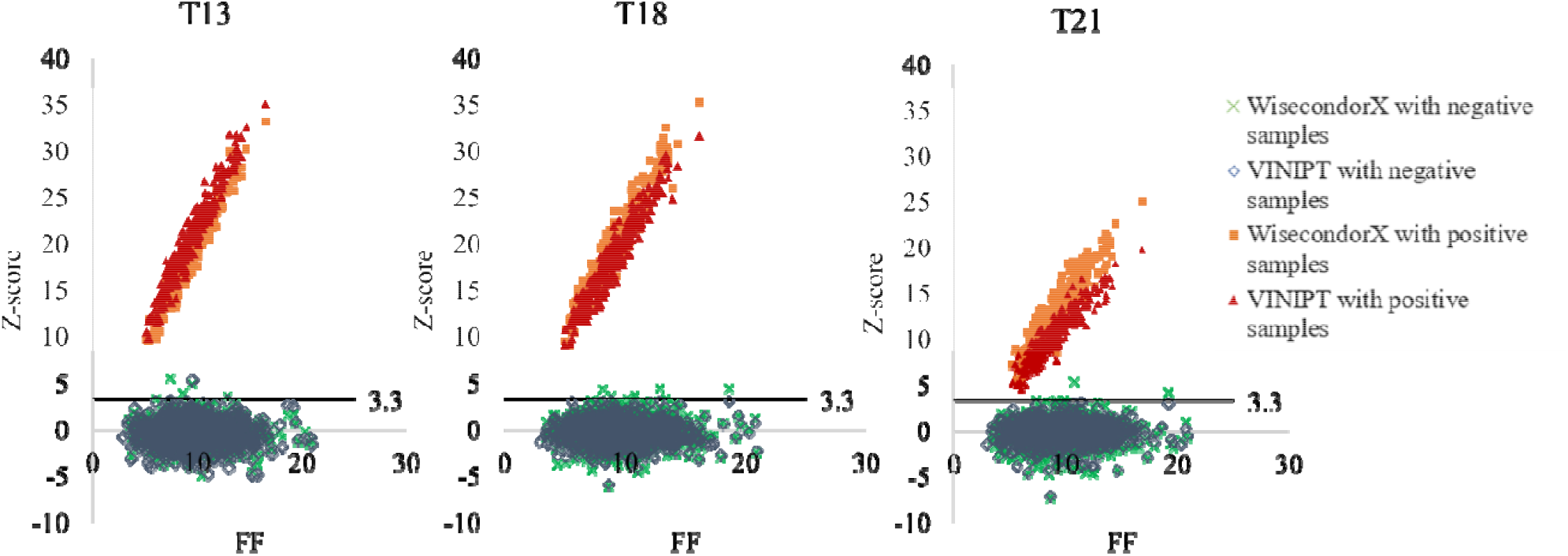
The z-scores of negative and positive non-mosaic samples. The left figure for T13, middle figure for T18, and right figure for T21. FF: fetal fraction

We now examined the results from WisecondorX and VINIPT for positive mosaic samples. It is much more difficult to detect aberrations from mosaic samples than from the non-mosaic samples. Fig. 3 shows that the z-scores of negative samples and that of positive mosaic samples are slightly overlapped.

**Fig 3.**
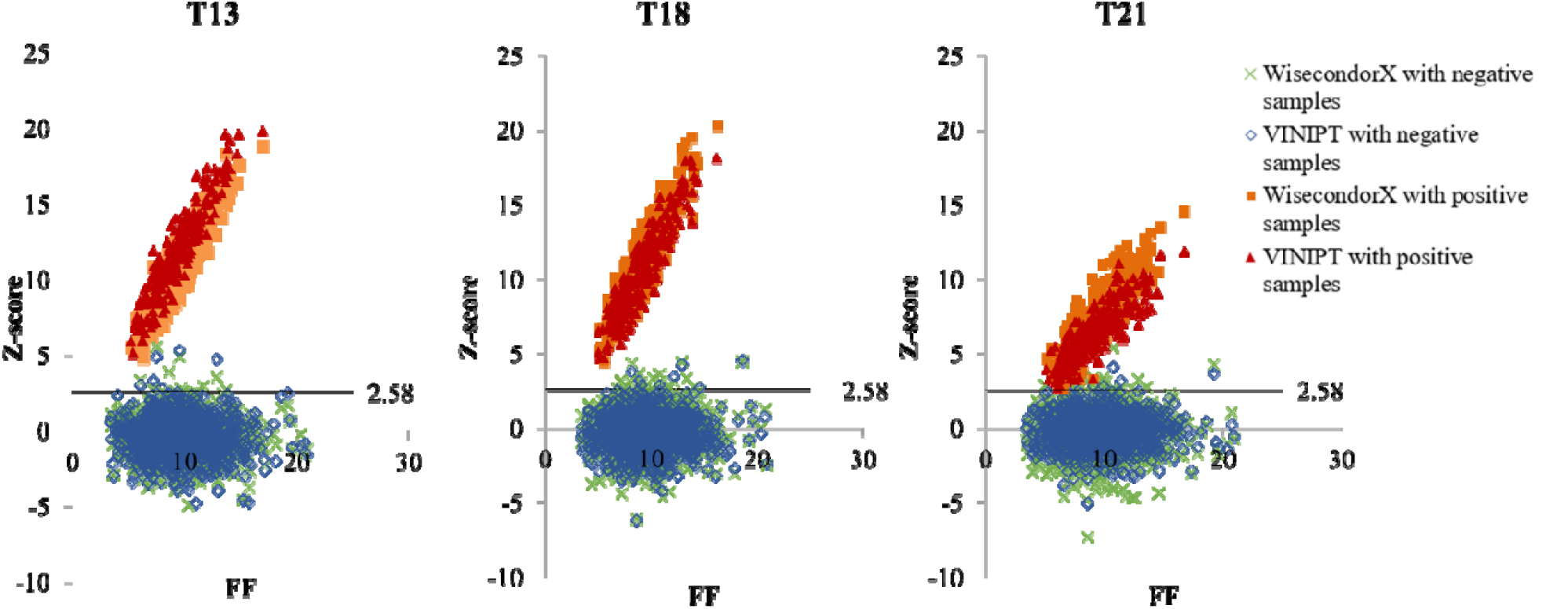
The z-scores of negative samples and positive mosaic samples. The left figure for T13, middle figure for T18, and right figure for T2. FF: fetal fraction

The z-score threshold of 3.3 resulted in a number of false negative results. To avoid a high false negative rate, we lowered the z-score threshold from 3.3 to 2.58 for both WisecondorX and VINIPT for the mosaic analysis. The WisecondorX and VINIPT were able to detect positive samples with T13 and T18 from the mosaic samples. Both WisecondorX and VINIPT algorithms falsely assigned two positive samples with T21 aberrations as negative samples. The overall sensitivity of detecting T13, T18, and T21 aberrations of the algorithms are 99.7%. The fetal DNA fractions of the two false negative samples are considerably low (5.8% and 6.1%). Lowering the z-score threshold helped reduce the false negative rate, however, it increased the false positive rate of both algorithms. WisecondorX resulted in 53 false positive samples, while VINIPT produced 19 false positive results. The false positive rates of WisecondorX and VINIPT for the mosaic analysis are about 5% and 1.8%, respectively.

Finally, we collected 100 positive samples with T13, T18, and T21 aberrations from our genetic testing center. They are called clinical positive samples to differentiate from the generated positive samples. TABLE 1 shows that the z-scores of the clinical positive samples and generated positive samples for both non-mosaic and mosaic conditions. The z-scores from generated positive non-mosaic samples are similar to that from clinical positive samples; and much higher than the z-scores from the generated positive mosaic samples. The results indicate that it is much more difficult for NIPT algorithms to detect chromosomal aberrations from mosaic samples than from non-mosaic

**TABLE 1.** THE Z-SCORE DISTRIBUTION OF 100 CLINICAL POSITIVE SAMPLES AND 1140 GENERATED POSITIVE SAMPLES WITH T13, T18 AND T21 ABERRATION.

|  | T13 | T18 | T21 |
| --- | --- | --- | --- |
| Clinical positive samples | 16.3 (8.5) | 20.8 (5.9) | 15.2 (6.8) |
| WisecondorX non-mosaicism | 17.9 (4.8) | 19.2 (5.2) | 12.7 (3.8) |
| VINIPT non-mosaicism | 19.5 (5.1) | 17.8 (4.7) | 10.7 (2.9) |
| WisecondorX mosaicism | 10.5 (2.7) | 11.2 (3.0) | 7.5 (2.3) |
| VINIPT mosaicism | 11.6 (2.9) | 10.3 (2.8) | 6.3 (1.7) |

## Discussion

The NIPT has been widely used as a powerful screening test to detect numerical abnormality of chromosomes. As the positive samples are rare, the positive datasets used to evaluate NIPT algorithms are typically small and not publicly available for scientific community. This prevents the development of new NIPT algorithms for different NIPT data types such as the low coverage NIPT data or the NIPT data obtained from mosaic samples.

A large number of negative NIPT samples are available from hospitals or genetic testing centers. In this paper, we present a computational method to create positive datasets from the negative samples. Our method does not introduce new short-reads; therefore, the created positive samples can play as real as the actual positive NIPT samples. We applied the method to create 1440 positive samples with T13, T18, and T21 aberrations for both non-mosaic and mosaic conditions.

We used our negative dataset together with the generated positive datasets to evaluate the performance of WisecondorX and VINIPT software. Experiments showed that both algorithms could determine almost all positive samples. They failed to detect a few positive mosaic samples with T21 aberration. We found that the false negative mosaic samples have considerably low fetal DNA fractions. The low amount of DNA from the fetus makes the NIPT algorithms more difficult to detect chromosomal aberrations from mosaic samples. For the mosaic analysis, we should use a lower z-score threshold; and require a high fetal DNA fraction to reduce the false negative rate.

The positive detection rate of NIPT, i.e., the percentage of cases reported positive by NIPT, ranges from 0.5% to 1.5% depending on a number of factors such as sequencing platforms, read coverage per sample and NIPT algorithms used to call aberrations. Different large-scale retrospective studies have showed that the positive predictive values of NIPT are considerable low. Therefore, positive results from NIPT must be carefully interpreted and explained by genetic consultants for pregnant women.

